# 4-methylumbelliferone enhances neuroplasticity in the central nervous system: potential oral treatment for SCI

**DOI:** 10.1101/2023.01.23.525137

**Authors:** Sîan F. Irvine, Sylvain Gigout, Kateřina Štěpánková, Noelia Martinez Varea, Lucia Machová Urdzíková, Pavla Jendelová, Jessica C. F. Kwok

**Affiliations:** School of Biomedical Sciences, Faculty of Biological Sciences, University of Leeds, Leeds, United Kingdom; Centre for Reconstructive Neuroscience, Institute of Experimental Medicine, Czech Academy of Science, Prague, Czech Republic; Department of Neuroscience, Charles University, Second Faculty of Medicine, 15006 Prague, Czech Republic

**Author notes:** These authors contributed equally to the manuscript (Sîan F. Irvine, Sylvain Gigout). These authors contributed equally to the manuscript (Lucia Machová Urdzíková, Pavla Jendelová, Jessica C. F. Kwok). Correspondence: Dr Jessica Kwok, School of Biomedical Sciences, Faculty of Biological Sciences, University of Leeds, Leeds, United Kingdom.

## Abstract

Perineuronal nets (PNNs) are specialised extracellular matrix (ECM) structures that act as key plasticity regulators to the central nervous system. Removal of PNNs using chondroitinase ABC injections restores plasticity, however, there are limitations to its application to due to its enzymatic nature. Here, we explore the use of a small molecule 4-methylumbelliferone (4-MU) as an alternative non-invasive strategy to reversibly remove PNNs and enhance plasticity. Oral administration of 4-MU for 10 days successfully and dynamically removed PNNs *in vitro*. While 4-MU, preferentially downregulated hyaluronan in the spinal cord, a down-regulation of chondroitin sulphate proteoglycans is also observed in the cortex. Long term administration for 8-weeks administration revealed a partial removal of PNNs, and that injury-induced mechanisms promoting cortical structural plasticity are linked to endogenous modulation of ECM molecules. 4-MU is a new tool to unravel the limits of normal and pathological PNN-mediated plasticity.

## 1 Introduction

Perineuronal nets (PNNs) are a compact and specialised form of extracellular matrix (ECM) uniquely found throughout the central nervous system (CNS) [1, 2]. The formation and presence of PNNs is strongly linked with the restriction of different forms of CNS plasticity, including the termination of developmental plasticity [3, 4]. PNNs have been implicated in the pathologies of various neurological disorders, including Alzheimer’s disease, epilepsy and schizophrenia [5–8], as well as in recovery from traumatic CNS injuries, such as spinal cord injury (SCI) [9, 10].

PNNs are complex and heterogenous structures composed of an assortment of proteoglycans and glycoproteins, assembled upon a skeleton of hyaluronan (HA) [11]. This provides a hierarchical condensation to a mesh-like network, where the maintenance or integrity of this structure is vital to its function [4, 12]. For example, the accumulation of inhibitory components in PNNs, including chondroitin sulphate proteoglycans (CSPGs) and sequestered proteins such as Sema3A [13], confers stability to synapses within the ‘holes’ of the net and protection against the formation of aberrant connections [14]. The removal of PNNs has been demonstrated to restore plasticity in multiple models throughout the CNS, including restoration of ocular dominance after dark rearing [3], erasure of fear memories [15] and compensation following incorrect repair of a peripheral nerve injury [16].

The most common methods for PNN removal harness enzymatic degradation of the constituent glycans in PNNs, particularly using the bacterial enzymes chondroitinase ABC (ChABC) and hyaluronidase. Of these, ChABC has been extensively used as an experimental tool to re-induce plasticity via the dual action of catabolising both chondroitin sulphate glycosaminoglycan (CS-GAG) chains and HA, in both the neural ECM and PNNs [16–19]. However, ChABC is an invasive therapy that requires intrathecal injections to the site of interest and due to its thermal instability requires repeated injections to sustain PNN removal [20]. Whilst much of the last few decades has focused on establishing means to maintain removal of PNNs by ChABC through the development of thermal stabilised and lenti-viral delivery systems [21–25], only recently have efforts been made to develop controllable method for PNN removal [26]. This latest iteration achieved this using a doxycycline-inducible regulatory transgene switch, which unfortunately still retained a low ‘leaky’ secretion of ChABC by mammalian cells without stimulus [26].

Whilst the development of this complex gene delivery system has allowed ChABC to maximise its potential, the prospect of a simple oral administration of a plasticity enhancer remains highly advantageous as both an investigative tool and a prospective therapy as it eliminates the associated ethical concerns and invasive risks. The following introduces a novel non-invasive and pharmacological compound that can control the duration of PNN removal, known as 4-methylumbelliferone (4-MU), as an investigative tool to reversibly remove PNNs and enhance plasticity throughout the CNS.

## 2 Results

The following results aim to investigate the efficacy of a non-invasive method of removing PNNs, using the compound 4-MU, in enhancing plasticity, using *in vitro* and *in vivo* methodologies. We aim to show that 4-MU can be used as an investigative tool in both short- and long-term studies *in vivo* and to demonstrate plasticity enhancement in a SCI model.

### 4-MU dynamically removes PNN components *in vitro* and *in vivo*

PNNs exist as a condensed lattice of various glycoproteins and proteoglycans, including HA and proteoglycan link proteins (HAPLNs) and the CSPG aggrecan (ACAN), bound and aggregated upon a HA backbone extruded from the cell surface [11, 27, 28]. A previously developed human embryonic kidney 293T (HEK) cell model, engineered to express the essential ECM components (hyaluronan synthase 3 (HAS-3) and HAPLN-1) required to induce the formation of a pericellular PNN-like structure (PNN^+ve^ HEK cells) [27], was used to investigate the efficacy of 4-MU in removing PNNs *in vitro* (Fig. 1A-B). Staining with the lectin *Wisteria floribunda* agglutinin (WFA), a sulphation-dependant ‘universal’ CSPG marker [12, 29], revealed normal ubiquitous ECM deposition by PNN^-ve^ HEK control cells (Fig. 1Ai). PNN^-ve^ HEK cells endogenously produce some ECM components, including ACAN [27], but lack a pericellular matrix coating (Fig. 1Ai). In contrast, untreated PNN^+ve^ HEK cells illustrated a clear upregulation of WFA binding, including PNN-like structures (Fig. 1Ai-ii). Two days of 4-MU treatment (0.5 mM or 1.0 mM) administered to PNN^+ve^ HEK cells removed 86.4 ± 4.47% (*F_3, 66_ = 73.60, p<0.0001 for all treated timepoints vs. untreated*) of WFA-positive staining which partially recovered to 43.6 ± 14.6% of baseline lectin binding within three days post-treatment (*p<0.0001 for 3 d and 5 d post-treatment vs. during*, Fig. 1Aiii-viii, B). These results indicate that 4-MU-mediated rem*oval of PNNs is dynamic and reversible*.

**Figure 1.**
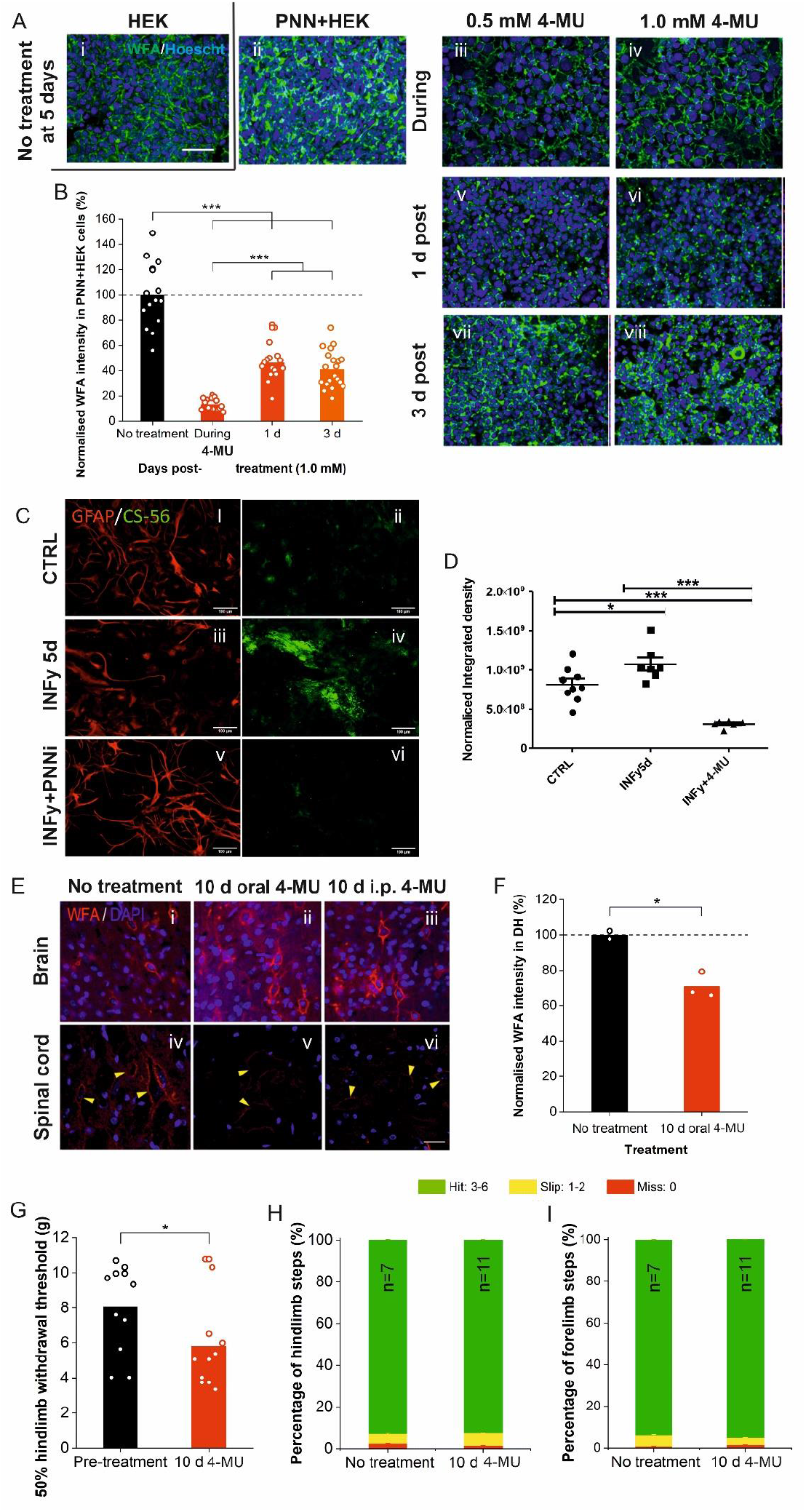

Reactive astrocytes are a main component of the glial scar that forms after CNS injury. Astrocytes (labelled using fibrillary acidic protein (GFAP)) show minimal secretion of CSPGs (labelled by CS-56; **Ci-ii, D**) until activated by five days treatment with interferon gamma (IFNγ; **Ciii-iv**). INFγ also produced morphological changes inducing protoplasmic morphology. Addition of 4-MU for 3 days to the activated astrocyte culture (INFγ+) resulted in downregulation of CS-56 staining compared to INFγ+ control **(Cv-vi, D)**. CS-56 reactivity in the INFγ+4-MU astrocytes was lower than in the unactivated astrocyte control, revealing an effective blockade of CSPGs secretion from astrocytes by 4-MU. This suggests that 4-MU has the potential to promote regeneration at the inhibitory glial scar after injury in vivo.

To determine if the small molecule 4-MU could induce PNN reduction in the CNS, we first perform a short-term 4-MU treatment *in vivo,* with administration (10 days) via either oral feeding or intraperitoneal (i.p.) injection twice daily. Histology from animals terminated after ten days dosing revealed that both methods of 4-MU administration were sufficient in decreasing WFA-positive binding throughout the CNS compared to non-treated animals (Figure 1J-O). Interestingly, 4-MU appeared to be more efficacious in downregulating the lectin binding in the spinal cord than in the cortex (Figure 1E). Importantly, this indicates that 4-MU, or its metabolites, can down-regulate the ECM in the CNS. Quantification in the dorsal horn of the spinal cord illustrated that ten days of oral 4-MU administration induced a partial removal of WFA-positive moieties to 71.0 ± 7.20% of baseline ECM levels (*t(3)* = *5.15, p*=*0.0142*; Fig. 1F). Short-term oral 4-MU also downregulated staining with HA binding protein to 64.0 ± 5.84 % (HABP; *t(3)* = *7.69, p*=*0.00456*, Supplemental Fig.1B) and for HAPLN-1 to 68.3 ± 8.09% (*t(3)* = *5.01, p*=*0.0153*, Supplemental Fig.1B) of baseline levels in the dorsal horn. As oral 4-MU treatment is both non-invasive and generated sufficient effects on the neural ECM *in vivo*, the following 4-MU dosing experiments with Lister Hooded rats used this method of administration.

### Short-term and partial removal of PNNs leads to hyperplasticity in uninjured spinal cord

Firstly, we questioned whether PNN removal throughout the CNS would affect normal sensation and motor functions. To investigate this, adult female Lister Hooded rats (*n*=*11*) were given ten days of oral 4-MU treatment and subjected to routine behavioural testing. These revealed that short-term 4-MU was sufficient to induce changes to sensory but not motor functions in the treated intact rats (Fig. 1G-I). Comparison to a pre-treatment baseline (7.9 ± 2.73 g) with the same rats, revealed that short-term 4-MU administration decreased the withdrawal threshold to 5.5 ± 2.03 g, representing an approximate 30% increase in sensitivity (*t(10)* = *2.76; p=0.02* Fig. 1G). Open field locomotor testing found no significant differences with short-term 4-MU treatment, with all animals achieving top scores of 21 on the Basso, Beattie, Bresnahan hindlimb (HL) scale [30] (*n.s.; p*=*1*). When a more skilled walking task was used to assess locomotion, HL locomotor activity also appeared consistent between 4-MU treated and untreated animals (Fig. 1H), with correct stepping (green) approximately 92.7 ± 0.82% (*n.s., t(16)* = *0.259, p*=*0.799*) of the time, with slips (yellow) and misses (red) making up 5.43 ± 0.70% (*n.s., t(16)* = −*0.978, p*=*0.343*) and 1.92 ± 0.41% (*n.s., t(16)* = *1.15*, *p*=*0.268*) of all steps, respectively, on the horizontal ladder. Similarly, forelimb (FL) performance was unaffected by short-term 4-MU (Fig. 1I), with 1.13 ± 0.24% missed (*n.s., t(16)* = −*1.64, p*=*0.121*), 4.35 ± 0.54% slipped (*n.s., t(16)* = *1.97, p*=*0.0667*) and 94.4 ± 0.56% correct steps (*n.s., t(16)* = −*1.34, p*=*0.197*). Regardless of treatment, HL and FL function performed equally as well with low percentages of stepping errors. Overall, acute removal of PNNs in the normal adult rat slightly increased sensitivity in the limbs tested, aligning with downregulation of PNN components in the dorsal horn with the same treatment paradigm, but did not affect locomotor function. No other behavioural differences were observed.

The sensorimotor cortex (M1) contains a highly organised topographical representation of motor movements that is subject to structural and functional plasticity in response to sensorimotor learning or neuronal injury [31–36]. Using intracortical microstimulation (ICMS), mapping of the HL and FL cortical movement representations were used to investigate the functional organisation of the M1 of intact/sham animals after long-term treatment with the prospective plasticity enhancer, 4-MU by comparing to the intact baseline HL and FL representations or ‘epicentres’ (see dotted lines; Fig. 2). Whilst 4-MU treatment reduced the total area able to elicit HL movements (*F_2, 12_* = *10.7, p*=*0.00764 for Sham/4-MU vs. Intact, p=0.00914 for Sham/4-MU vs. Sham*), this was due to some of the intact HL epicentre being no longer able to elicit HL movements (*F_2, 12_* = *18.6*, *p*=*0.00411 for Sham vs. Sham/4-MU, p<0.001 for Intact vs. Sham/4-MU*; Fig. 2A-H). Paired-pulse protocols [37, 38] were used as an index to study possible alteration of intracortical GABAergic inhibition in parallel, using *in vitro* sensorimotor cortical slices, where the paired-pulse ratios (PPR) for short and long interstimulus intervals allowed us to investigate GABA_A_ and GABA_B_ receptor-mediated inhibition, respectively. Healthy control slices exhibited a pronounced paired-pulse inhibition (PPR: ~0.5), however, this was not altered with either sham surgery or 4-MU treatment (*n.s., F_2,65_* = *1.52, p*=*0.227*, Fig. 2I; *F_2,65_* = *2.35, p*=*0.0960*, Fig. 2J). This suggested that the observed 4-MU-induced decreases in HL area was due to structural reorganisation and not alteration of cortical inhibition. Like that observed with the HL representation, 4-MU induced functional reorganisation of the FL representation in sham animals (Fig. 2K-S). Despite no change in the total surface area that elicited FL movements (n.s., *F_2, 12_* = *0.249, p*=*0.784*; Fig. 2N), the incidence of these were reduced in the intact FL epicentre (*F_2, 12_* = *5.61, p*=*0.0226*; Fig. 2P) but did not appear to encroach on the HL area (*n.s., F_2, 12_* = *0.900, p*=*0.437*, Fig. 2R; *F_2, 12_* = *2.38, p*=*0.142*, Fig. 2S). There was no significant change in cortical stimulation thresholds required to elicit HL or FL movements with 4-MU treatment (Supplementary Table 1). As short-term 4-MU administration increased tactile sensitivity and long-term 4-MU treatment induced structural reorganisation, we can conclude that the partial removal of PNNs by 4-MU, as illustrated above, was sufficient in producing an environment favourable to promoting functional plasticity at multiple levels of the CNS. As these plastic changes have been observed in treated intact animals, this may result in maladaptive connectivity.

**Figure 2.**
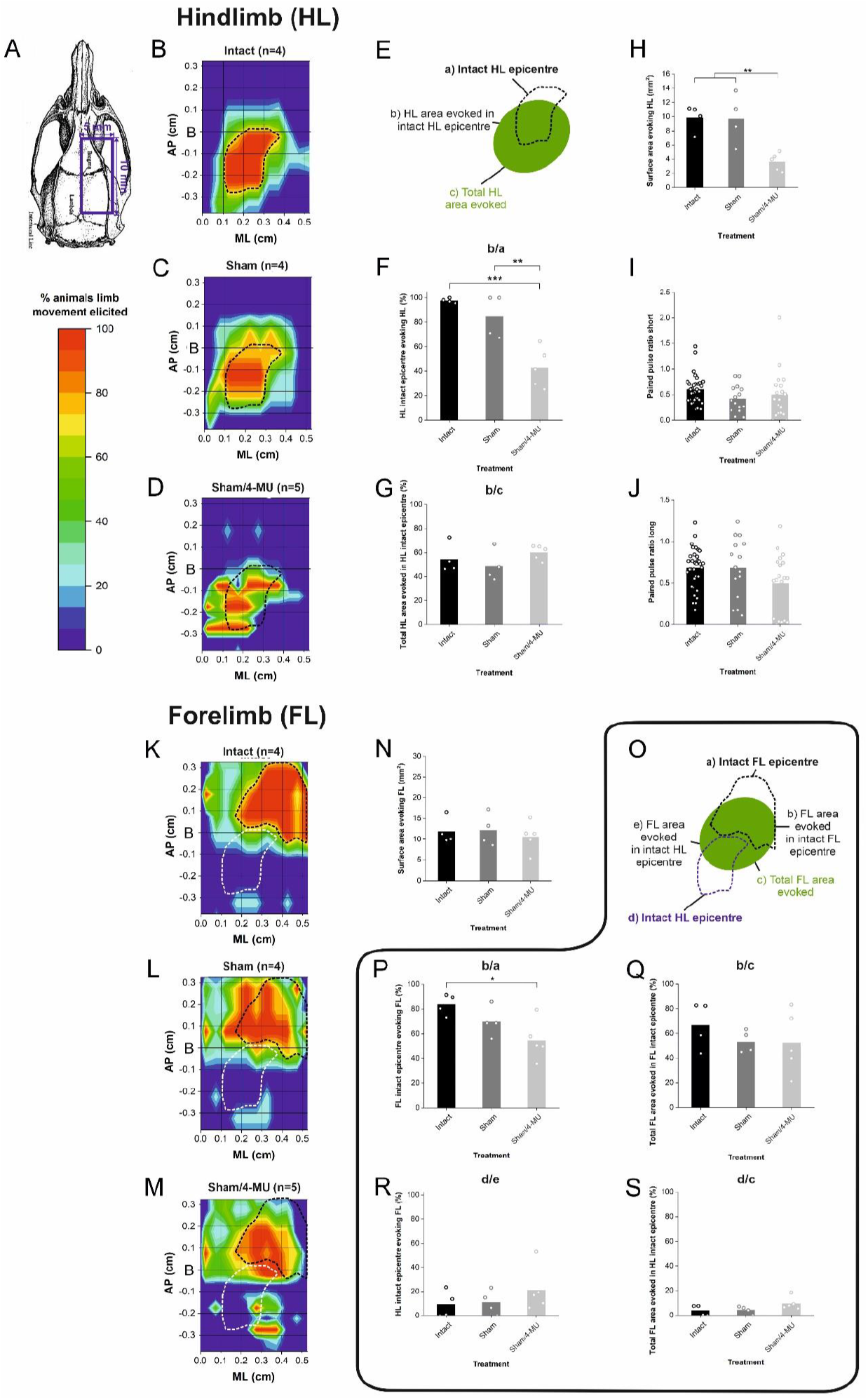

### 4-MU induces differential effects between the brain and spinal cord in removing PNN components

The initial findings above indicated that 4-MU may have a stronger effect in reducing PNN components in the ECM, labelled by WFA, in the spinal cord than in the brain following short-term treatment in intact rats (Fig. 1E). Further investigation with long-term (12 weeks) 4-MU treatment indicated that whilst in the brain overall GAG content was lowered by approximately 60% by 4-MU (2.65 ± 0.67 to 1.01 ± 0.35 mg/g wet weight brain, *t(10)* = *5.28, p*<*0.001*, Fig. 3B), the picture was more complex and various PNN components were differentially regulated in distinct CNS regions (Fig. 3). In the M1, immunohistochemical labelling for PNN components identified PNNs throughout layers II-VI and especially clustered in layers V/VI (Fig. 3F-K). 4-MU treatment in sham animals downregulated both ACAN- and WFA-positive staining by almost 60% (ACAN: 41.9 ± 6.72%, *t(4)* = *3.91*; *p*=*0.0477*; WFA: 43.1 ± 11.0%, *t(4)* = *3.80*, p= 0.0191; Fig. 3C-K), highlighting a decrease in CSPGs in the cortical ECM. However, hyaluronan binding protein (HABP) intensity displayed a modest yet not significant decrease in HABP binding in the cortical ECM to 64.0 ± 32.0% of baseline levels following 4-MU treatment *(t(4)* = *1.67, p*=*0.170*, Fig. 3D). The spinal cord exhibited the inverse trend, where 4-MU treatment significantly downregulated HA, denoted by HABP, in the ventral horn (VH) ECM *(t(4)* = *6.78, p*=*0.00248*), yet CSPG content of the ECM, as labelled by ACAN and WFA, were unchanged from baseline levels (*n.s*., ACAN: *t(4)* = *1.05, p*=*0.354*; WFA: *t(4)* = *1.22, p*=*0.288*; Fig. 3L-S, Supplemental Fig. 2).

**Figure 3.**
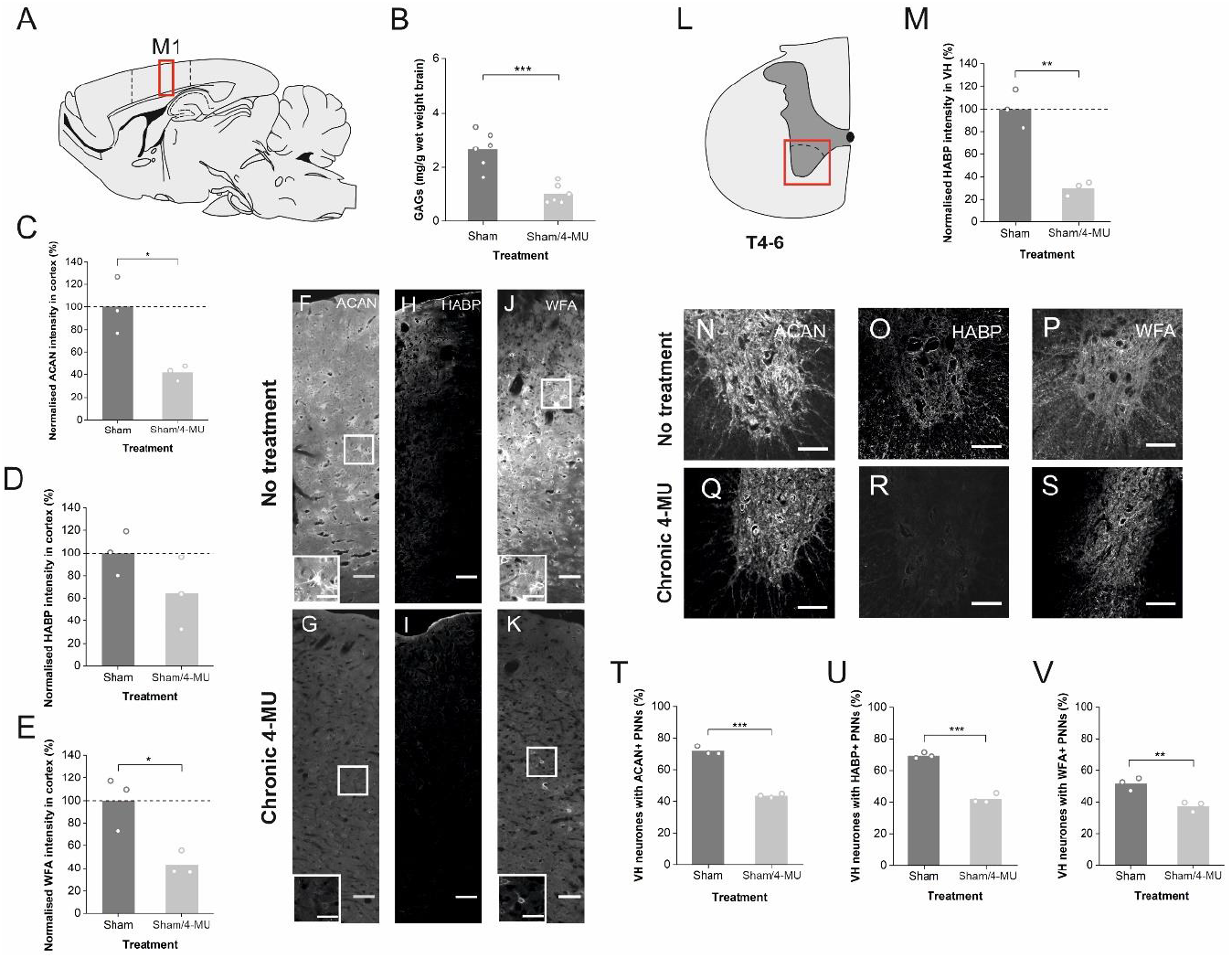

Despite the lack of significant influence of 4-MU treatment on CSPG content in the spinal ECM, a significant yet partial decrease in the number of PNNs surrounding VH neurones was observed (Fig. 3T-V). A similar baseline and decrease in PNN number was observed for both ACAN- (from 71.7 ± 2.71% to 43.5 ± 1.23%, *t(4)* = *16.4, p*<*0.001*, Fig. 3T) and HABP-positive PNNs (from 69.2 ± 1.82% to 42.1 ± 3.06%, *t(4)* = *13.2*, *p*<*0.001*, Fig. 3U), with fewer baseline PNNs and a more minor reduction of WFA-positive PNNs (from 51.3 ± 3.91% to 37.4 ± 2.00%, *t(4)* = *4.63*, *p*=*0.00983*; Fig. 3V).

Contrary to the effects illustrated with short-term 4-MU administration, long-term administration appears to have had an accumulative effect, revealing differential downregulation of various PNN components in distinct CNS regions. Additionally, the strong removal of HA in the cord appears to reflect the partial removal to a consistent number of PNNs, regardless of ECM labelling. Long-term 4-MU treatment partially removed PNNs, indicating that this is sufficient to promote plasticity, as illustrated by the motor map reorganisation above.

### 4-MU and/or injury independently enhances cortical plasticity in spinal cord injured rats

The prior experiments have all applied 4-MU to intact *in vivo* systems. We then performed the investigation on the effect of 4-MU in SCI, with long-term treatment to adult female Lister Hooded rats after a mid-thoracic moderate contusive SCI. Echoing the trend observed above, 4-MU treating injured animals removed GAGs from the brain by approximately 50% (1.26 ± 0.35 mg/g wet weight brain, *F_2,17_* = *17.6, p*<*0.001*, Fig. 4E), but these were preferentially CSPGs (ACAN: 37.3 ± 30.1%, *F_2,8_* = *8.18; p*=*0.0316 for SCI/4-MU vs. Sham*, Fig. 4B; WFA: 46.1 ± 15.3%, *F_2,8_* = *11.4, p*=*0.0243 for SCI/4-MU vs. Sham*, Fig. 4D), rather than HA (*F_2,8_* = *3.07, p*=*0.145* in Fig. 4C; Fig. 4A-F) and the inverse was observed in the spinal cord (HABP: *F_2,8_* = *22.8, p*=*0.0122 for SCI/4-MU vs. SCI/Vehicle, p=0.00174 vs. Sham; n.s*., ACAN: *F_2,8_* = *2.88, p*=*0.133*; WFA: *F_2,8_* = *3.38, p*=*0.104*; Fig. 4G-I, Supplemental Fig. 2). However, in the brain, injury also independently caused a comparable profile of GAG content to that of 4-MU treatment (1.09 ± 0.32 mg/g wet weight brain, *p*<*0.001*, Fig. 4E); in particular, after SCI there was a similar reduced CSPG content in the M1 (ACAN: 37.8 ± 7.52%, *p*=*0.0438 for SCI/Vehicle vs. Sham*; WFA: 39.7 ± 8.64%, *p*=*0.0144 for SCI/Vehicle vs. Sham*, Fig. 4A-F). In contrast, the injured spinal cord displayed partial reduction in PNNs (HABP: 36.7 ± 3.17%, *F_2, 9_* =*71.0, p*<*0.001*, Fig. 4K), including those labelled by CSPG markers (ACAN: 39.6 ± 3.67%, *F_2, 9_* =*141.7*, *p*<*0.0001*, Fig. 4J; WFA: 35.1 ± 4.07%, *F_2, 9_* =*31.5, p*<*0.001*, Fig. 4L), only after 4-MU treatment (Fig. 4G-L).

**Figure 4.**
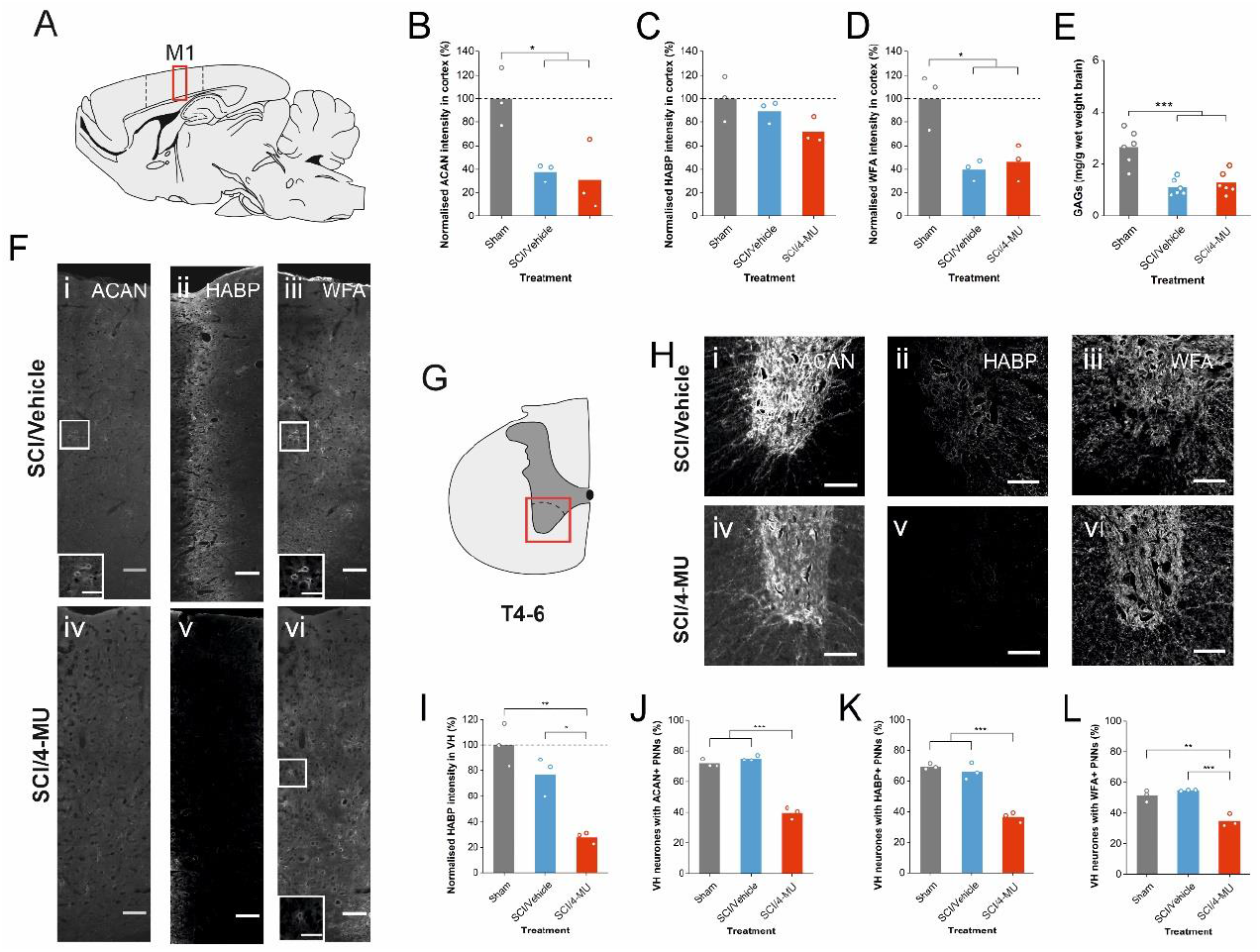

After mid-thoracic SCI, HL movements were unable to be elicited by ICMS (Fig. 5B-C), despite evidence of open-field HL ambulation, whereas FL movements were able to be elicited in both intact FL and HL areas (Fig. 5D-E). Injury and/or treatment caused no changes to the minimum or mean cortical stimulation thresholds required to elicit FL movements (Supplemental Table 1). There was no total change in the surface area eliciting FL movements with injury and/or long-term 4-MU treatment (*n.s., F_2,9_* = *0.403, p*=*0.683*, Fig. 5F), nor the area associated with the intact FL epicentre (*n.s., F_2,9_* = *0.891, p*=*0.452*, Fig. 5K; *F_2,9_* = *0.0685, p*=*0.934*, Fig. 5L). However, the combination of both injury and long-term 4-MU treatment caused a larger percentage of FL movements (from 4.27 ± 1.56% to 23.9 ± 8.01%) to be associated with areas previously eliciting HL movements, indicating an overall shift, not expansion, of the group’s representative FL area into the intact HL epicentre, compared to sham control (*F_2, 9_* = *2.58, p*=*0.197 for Sham vs. SCI/4-MU*, Fig. 5M; *F_2, 9_* = *5.23, p*=*0.0432 for Sham vs. SCI/4-MU*, Fig. 5N).

**Figure 5.**
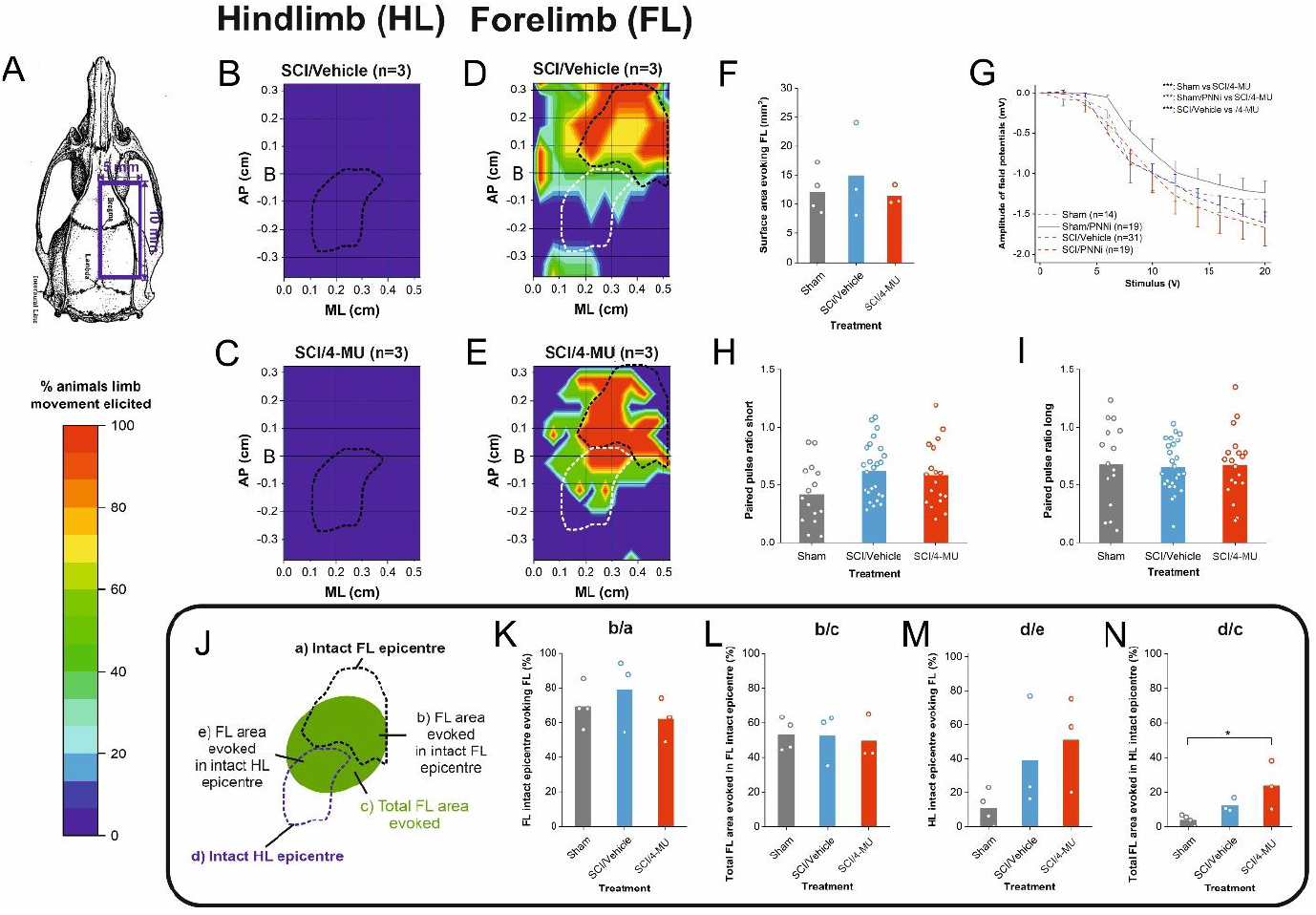

Parallel experiments addressing the local excitability of *in vitro* cortical slices, used individual input-output curves of field potential amplitudes (Fig. 5G) which were fitted by the Boltzmann equation, yielding the PSP_max_, the I_50_ and the slope factor (Supplemental Table 2). Importantly, this revealed that the combination of SCI and 4-MU treatment gave results significantly different from all other experimental groups (*Greenhouse-Geisser F_1.49, 13.4_* = *1.61, p*<*0.001 for all groups vs. SCI/4-MU*, Fig. 5G), suggesting that together 4-MU and injury created an environment of increased intracortical synaptic transmission, particularly at higher stimulation intensities, which may contribute to the reorganisation of the M1 observed with this group. However, further analysis revealed an increased slope factor after SCI, indicating decreased efficacy of the intracortical system (*F_4, 108_* = *6.85, p*=*0.00529 for SCI/Vehicle vs. Intact, p=0.0230 vs. Sham, p<0.001 vs. Sham/4-MU, p=0.0196 for SCI/4-MU vs. Sham/4-MU*; Supplemental Table 2). Addition of extracellular 4-MU in the artificial cerebral spinal fluid (ACSF) ([4-MU]_e_) increased the PSPmax in each group by approximately 20%, indicating an increase in intracortical synaptic response (Supplemental Table 3). These effects were independent of prior conditions (SCI or long-term 4-MU treatment), suggesting that the differences in intracortical inhibitory/excitatory balance observed between groups is likely due to the effects of injury and long-term oral 4-MU treatment (*n.s. for all parameters*, Supplemental Table 3). Therefore, the off-target (not PNN/ECM) effects of 4-MU does not appear to have been down-regulated or desensitised with long-term oral 4-MU treatment, as the increase in PSP_max_ by [4-MU]_e_ was also observed in slices from Sham/4-MU and SCI/4-MU animals.

Paired-pulse protocol also revealed that injury alone, not the SCI/4-MU combination, affected paired pulse responses at short but not long interstimulus intervals, with an increase in PPR from 0.42 ± 0.26 to 0.63 ± 0.24 (*F_2, 57_* = *3.09, p*=*0.052 for Sham vs. SCI/Vehicle*, Fig. 5H-I), implying that SCI induced a decrease in GABAA-receptor mediated transmission. In summary, despite the ability of both injury and 4-MU to independently induce similar decreases in CSPG content in the M1, only long-term 4-MU administration could enhance plasticity sufficiently enough to foster reorganisation of the cortical motor map, suggesting that additional mechanisms, such as the observed increase in local intracortical excitability, may underly these functional changes.

## 1 Discussion

PNNs can be removed through either degradation of existing PNNs or via interference with the dynamic formation or ‘turnover’ of the PNNs. Whilst enzymes including ChABC have been liberally utilised in PNN research, we have presented here a novel pharmacological agent as an alternative and non-invasive method of investigating the plastic effects mediated by PNN removal. Using *in vitro* and *in vivo* methods, we demonstrated that 4-MU differentially downregulates key 4-MU components in the ECM and was able to remove PNNs at multiple levels of the CNS. Importantly, 4-MU induced functional plasticity after both short- and long-term administration, although this could be maladaptive in intact systems. It was established that removal of some but not all PNNs was adequate to restore plasticity to both the intact and injured CNS.

As PNNs are a dynamic and heterogeneous structure, the turnover of its components affects the overall integrity of the architecture. In the spinal cord, 4-MU appeared to have produced a strong but incomplete downregulation in HA production (~25% remaining) in both the loose ECM and in PNNs. Whilst other PNN components were found to be distributed in normal concentrations in the loose ECM, the absence of the HA pericellular coat - the backbone of the PNN structure [11], prevented the aggregation of the other key components, including CSPGs. In the brain, 4-MU appeared to alter PNNs instead by decreasing the overall CSPG expression in the ECM and therefore the aggregation of the highly inhibitory CSPGs into the PNN structure. Whilst it is clear that 4-MU has exerted differential effects on the brain and spinal cord, short-term 4-MU administration suggested that a stronger and more specific PNN effect was detected in the cord than in the brain, correlating with growing evidence that PNNs in the brain and spinal cord differ in composition. However, long-term administration exhibited a more powerful and possibly accumulative effect on the brain. Further research is required to elucidate the discrepancy between these effects and to advance understanding of the mechanism of 4-MU in the CNS.

Although 4-MU and ChABC both aim to remove PNNs to enhance plasticity, injection of ChABC into the CNS results in a broad and non-specific degradation of CS-GAGs and HA in both the diffuse ECM and in PNNs [18, 19], whereas the effects of 4-MU seem to be much more specific to PNNs and its effects on inhibitory CSPG signalling in the ECM are region specific [39–41]. Crucially, 4-MU demonstrated that specifically removing some but not all PNNs throughout the CNS can promote plasticity. In contrast, ChABC-mediated plasticity is limited to the local digestion area, whereas 4-MU renders plasticity throughout the system. Theoretically, removal of PNNs from a larger area of the CNS should provide a greater scope for plasticity and thus correlate with superior functional recovery after CNS injury, for example. This is supported, albeit on a smaller scale than mediated by 4-MU, by the comparison of single dose bacterial ChABC injection versus lenti-viral ChABC, which digested a greater area proximal to the lesion as well as conferring enhanced functional recovery [24]. Whilst some conditional CNS knockouts of PNN components do exist, including tenascin-R, HAPLN-1 and ACAN [4, 42–45], and have proven ability to enhance plasticity, few of these have explored the effects in neuronal injury models or at multiple CNS levels [4, 43]. Of these only HAPLN-1 knockout animals have been applied to CNS injury models and unfortunately the effects exhibited in this study are limited to assessments of sprouting and not to functional changes [4]. Whilst not clinically relevant, applying a CNS injury to these conditional knockout models may help to elucidate the varying recovery profiles between partial and complete removal of PNNs throughout the CNS and help define the limits of PNN-mediated plasticity.

We have concluded that long-term administration of 4-MU removes some but not all PNNs in the investigated regions, yet it is unknown whether this is linked to the function or heterogeneity of these PNN populations and their associated neurones. Spinal PNN-associated neurones are known to temporally express different isoforms of the enzyme responsible for HA expression, HAS [11]. It is possible that 4-MU affects different HAS isoforms, resulting in the observed sensitive and insensitive PNN populations to 4-MU-mediated removal, as well as the production of different lengths of HA chains [11]. However, more research is required into the mechanism of 4-MU to elucidate this. Importantly, this highlights that the development and use of a diverse range of investigative tools is beneficial to probe out different aspects of the function and structure of PNNs. For example, this study has already revealed that the injury-induced mechanisms promoting structural plasticity in the M1 are linked to endogenous modulation of the ECM molecules, akin to the decrease in CSPG concentration induced by 4-MU. Other studies concur that SCI is able to modulate the effect of PNNs in the M1 cortex, with downregulation of PNN thickness but not total number, as well as the inhibitory CSPG receptor NgR, particularly in the transition areas between FL and HL maps [46, 47].

The widespread nature of 4-MU in promoting plasticity throughout the CNS could be applied to PNN-related pathologies, such as improving memory retention [48, 49], or fostering recovery after traumatic brain injury or stroke [50–53]. In neuronal injury, particularly SCI, a more widespread plasticity enhancement could promote alternate connections at multiple spinal levels to improve functional recovery. The differing efficacy and ECM targets between the brain and spinal cord, particularly in short-term treatment, could prove beneficial to brain-related pathologies where more subtle modulation of these larger networks is required. The heterogeneous structure of PNNs has proved a challenging target to link to its dynamic properties and function in the CNS. The development of a diverse and dynamic toolkit is essential to advance our understanding of the limits of its role in regulating plasticity. Our evidence presents 4-MU as an alternative novel and non-invasive tool to investigate both the normal and pathological function of PNNs.

## 3 Methods

### 1.1: In vitro

#### 1.2: Animals

Adult female Lister-hooded rats (200-250 g; n=71) were obtained from Charles River Laboratories (Canterbury, UK) and housed in pairs in Central Biomedical Services (University of Leeds, UK) in a temperature-controlled environment in (20 ± 1 °C), with a 12 hr light/dark cycle (lights on at 07:00). Access to food and water was *ad libitum*. All procedures and experiments complied with the UK Animals (Scientific Procedures) Act 1986, following the 3R principles and the ARRIVE guidelines.

### 1.3: *In vivo* 4-MU treatment

To investigate the short term effect of 4-MU and to optimise the method of administration, adult female rats received 10 days administration of x dose 4-MU *i.p*, oral 4-MU (2 g/kg from a stock solution of 0.2 g/ml; n=x Sprague Dawley and n=11 Lister Hooded) or no treatment (n=7 Lister Hooded). Carboxymethyl cellulose (CMC; Sigma; 0.4%) was used to (partially) solubilise 4-MU (0.1 g/kg) for i.p injection.

Pharmacological treatment (4-MU; 2 g/kg from a stock solution of 0.2 g/ml) commenced from the day of injury/surgery for Lister Hooded rats (n=12 Sham/4-MU animals and n=11 SCI/4-MU). Following *in vitro* and short-term *in vivo* results, long-term 4-MU administration consisted of twice daily oral 4-MU via syringe (5 d/week for 12 weeks).

#### 1.4: Surgery

Using isofluorane, a T8 dorsal laminectomy was performed on Lister Hooded rats, followed by a 200 kdyn contusion injury (moderate-severe; *n*=*23*) [54] (Infinite Horizon impactor, Precision Systems and Instrumentation, LLC, Fairfax Station, VA) at the exposed T9 cord. Sham-operated animals (*n*=*23*) underwent the laminectomy but not the contusion injury. Analgesia (Vetergesic Buprenorphine; 0.015 mg/kg; Henry Schein, UK) and antibiotics (Baytril enrofloxacin; 2.5 mg/kg; Henry Schein, UK) were delivered subcutaneously for prescribed duration. Temporary individual housing allowed a week of recovery during this period of reduced mobility. Post-injury, bladders were expressed twice daily until urinary reflexes recovered (~2-3 weeks post-injury). All animals were monitored daily until the end of the experiment. The magnitude of the contusion injury was sufficient to impair locomotor and ambulatory ability in the injured rats only. All animals thrived with no animals required to be killed for health reasons.

#### 1.5: Experimental design

Intact Lister Hooded rats (n=7 no treatment; n=11 10 d oral 4-MU) underwent behavioural assessments after treatment before termination. For all behavioural tests, animals were assessed in chronological order independent of treatment group.

Lister Hooded rats (Sham n=23, SCI n=23) were received 12 weeks of 4-MU or no treatment/Vehicle. At 12 weeks animals were used for intracortical microstimulation (ICMS: Intact n=4, Sham n=4, Sham/4-MU n=4, SCI/Vehicle n=3, SCI/4-MU n=3), *in vitro* electrophysiology (Intact n=10, Sham n=4, Sham/4-MU n=5, SCI/Vehicle n=6, SCI/4-MU n=5) or GAG extraction (n=3 per group). For histology following long-term 4-MU treatment, following ICMS animals were terminated and transcardially perfused for use in histology (n=3 per group).

All images were randomised, and treatment groups were blinded to the analyser.

#### 1.6: Behavioural functional assessments

To determine the functional effects of acute 4-MU treatment, behavioural assessments were performed on intact Lister Hooded rats only, receiving no treatment or 10 d oral 4-MU.

##### 1.6.1: Basso, Beattie and Bresnahan HL locomotor open field test

HL locomotor ability was assessed using the Basso, Beattie and Bresnahan (BBB) HL locomotor scale [30]. BBB testing (4 mins/animal) was carried out in the morning using an open locomotor field (custom-built Perspex O-ring: diameter 80 cm, height 30 cm). Each BBB test was simultaneously assessed by two individuals to minimise subjective biases.

##### 1.6.2: Horizontal ladder

Rats were familiarised on a ‘training’ arrangement of rungs and tested on a ‘trial’ arrangement of rungs filmed using a camera (five trials; Canon Powershot SX720 HS; 30 frames/second at 10x optical zoom; 1.5 m distance from Plexiglas walkway). Trials were analysed frame by frame assessing each gait cycle for foot faults and paw placement, as described by Metz and Whishaw [55]. Steps were categorised as either a miss (score: 0), slip (score: 1 or 2) or hit (score: 3-6) and normalised by number of steps per trial.

##### 1.6.3: Von Frey assay

This assay evaluated changes to HL sensory function. Animals were acclimatised to Perspex cages with a wire mesh bottom (Complete Base assembly for plantar stimulation; Ugo Basile) immediately prior to testing until general movement and grooming activities stopped. Von Frey filaments (Touch Test™ Sensory Evaluator Kit of 20; #39337500; Leica Biosystems, England) were depressed against the plantar arch of the HL footpads. Sensory testing procedure and analysis was carried out as described by Chaplin et al. (1994) [56] using the Dixon up-down method [57] to determine the 50% withdrawal threshold for each HL.

#### 1.7: Electrophysiology

##### 1.7.1: *In vivo* intracortical microstimulation

Anaesthesia was induced using an i.p. injection of ketamine/xylazine (Ketavet®, Henry Schein, 100 mg/kg; Rompun®, Henry Schein, 3.2 mg/kg) and a jugular cannulation provided anaesthesia maintenance (10-20 mg/kg ketamine) to preserve cortically evoked motor responses. A craniotomy on the right hemisphere exposed the entire motor cortex (5 mm posterior to 4 mm anterior and 5 mm lateral, relative to bregma), keeping the dura intact. Time taken from induction to the full exposure of the above described motor cortex area allowed for adequate reduction of the depressant effect of xylazine on cerebral cortex neurones to perform intracortical microstimulation (ICMS) [58, 59].

ICMS was used to investigate functional reorganisation of the motor map. Using bregma as a cranial landmark, randomised electrode penetrations at stereotactic coordinates were made perpendicular to the pial surface (depth 1.3–1.7 mm) using a monopolar tungsten electrode (WE30030.5A5; Microprobes for Life Science, Maryland, USA; 0.2 ms pulses at 333 Hz for 40 ms, interval of 3 s between stimulation trains). The exploration grid covered the entire area of the motor cortex using at least 70 stimulation points with a minimum distance of 500 μm between points). Sites were randomly stimulated to minimise biasing from stimulation-evoked alterations to cortical representations [60, 61] or anaesthesia level.

Successful stimulation evoked a motor response with a current <40 μA to prevent wide-spread cortical activation. For each positive stimulation site, the minimum current threshold for each evoked response was recorded alongside the side (contralateral/ipsilateral) and type of response (FL/HL) and the muscles stimulated. HL and FL movements were evoked almost exclusively on the contralateral side of the body. Individual current activation thresholds were defined independently when more than one type of movement was observed at a single stimulation site.

Following completion of the ICMS protocol, animals were terminated, as described below, to perform histological assessment of PNNs.

##### 1.7.2: *In vitro* electrophysiology

Sham or injured rats (n= 9 or 11, respectively) received an overdose of isoflurane before neck dislocation. Coronal slices (400 μm; containing the sensorimotor cortex/M1) were made using a vibratome (VT100S; Leica Microsystems, Buffalo Grove, IL, USA) in cold oxygenated (carbogen: 95% O_2_ – 5% CO_2_) artificial cerebral spinal fluid (ACSF; in mM, 124 NaCl, 26 NaHCO_3_, 1.25 NaH_2_PO_4_, 3 KCl, 1.5 CaCl_2_, 1.5 mM MgCl_2_, 10 glucose; pH 7.4), left for recovery (RT; 2 hrs) before incubation at the recording chamber between humidified carbogen gas and oxygenated ACSF (33.0 °C).

Synaptic responses were elicited by electrical stimuli (0–20 V, increment 2 V, 100 μs duration at 0.1 Hz, in triplicate) applied via a bipolar tungsten electrode placed in the deeper cortical layers (V/VI), with responses recorded locally (2–4 MΩ recording electrodes; ACSF filled). Signals obtained were fed to Axoclamp 2B amplifier (Molecular Devices, Sunnyvale CA, USA). Recorded signals were digitised online (10 kHz) with a PC based system (Digidata 1322A and Clampex 10.3 software, Molecular Devices, Sunnyvale CA, USA) and analysed off-line (Clampfit 10.3).

For each slice the average field potential amplitude was plotted versus the stimulus intensity. Then on each of these input-output curves, a Boltzmann fit was performed, and three key values were extracted (maximal amplitude of the field potential (PSP_max_), the stimulus yielding half-maximal response (I50) and the slope factor) and the average value per slice was grouped. 4-MU was dissolved in dimethylsulfoxide (DMSO) before being added to ACSF to a final concentration of 500 μM 4-MU and <0.1% DMSO. 4-MU (500 μM) was added to ACSF immediately prior to recording and for at least 30 mins. We checked that responses were abolished by kynurenic acid (1 mM) at the end of experiments [62], confirming that they were synaptic.

Paired-pulse protocol was performed to evaluate the possible effects of 4-MU on neurotransmitter release via presynaptic sites.

The paired pulse ratio (PPR) was calculated for several interstimulus intervals (10 ms - 1 s) using the “amplitude of the second field potential/ amplitude of the first response” to identical stimulations (100 μs, 20 V). 20 V was chosen because it induced a maximal response with a pronounced paired-pulse depression. We defined a short (20-40 ms) and a long (150-250 ms) interstimulus interval which coincide respectively to the peak of GABA_A_ and GABA_B_ receptor-mediated responses. Both long and short PPR procotols were performed on the same slices. For each cortical slice, the average of three intervals for the short (20-40 ms with 10 ms step) and long (150-250 ms with 50 ms step) interstimulus intervals were calculated.

#### 1.8: Histology

##### 1.8.1: Tissue preparation

Animals were given an overdose of sodium pentobarbital (Pentoject; Henry Schein; 200 mg/kg; i.p. injection), before a transcardial perfusion (Gage, Kipke and Shain 2012) was performed using sodium phosphate buffer (PB; in M, 0.12 NaH_2_PO_4_; 0.1 NaOH; pH 7.4) followed by 4% paraformaldehyde (PFA; in PB; pH 7.4) for tissue fixation. The CNS was dissected, post-fixed in PFA (4%; 4 °C) overnight and cryoprotected in 30% sucrose solution (30% v/w sucrose in PB; 4 °C) until tissue saturation. The appropriate CNS segments were excised and frozen in optimum temperature medium (OCT; Leica FSC 22 Frozen Section Media; Leica Biosystems) to section (40 μm free-floating) using a cryostat (Leica CM1850; Leica Biosystems). Sections were serially collected into physiological buffer solution (PBS; in M, 0.13 NaCl, 0.7 Na_2_HPO_4_, 0.003 NaH_2_PO_4_; pH 7.4) to remove the OCT and stored in 30% sucrose solution (4 °C).

##### 1.8.2: Immunohistochemistry

At room temperature (RT), sections were washed (3 x 5 min) in Tris-buffered saline (TBS; in M, 0.1 Tris base, 0.15 NaCl; pH 7.4) to remove sucrose residue. Tissue was then blocked in 0.3% TBST (1× TBS solution and 0.3% v/v Triton X-100) and 3% normal donkey serum (NDS; v/v) for two hours before incubation in blocking buffer (3% NDS in 0.3% TBST; pH 7.4) containing primary antibodies (24 hrs; 4 °C; Table 2).

**Table 1:** Immunohistochemical detection of extracellular matrix components and neuronal markers, including concentration (conc.) of antibody.

<sup>1</sup> DSHB, Developmental Studies Hybridoma Bank, University of Iowa, USA. <sup>2</sup>WB, Western blotting
| Detected component | Marker | Host | Antibody Conc. | Source | Characterisation |
| --- | --- | --- | --- | --- | --- |
| **Extracellular matrix components** |  |  |  |  |  |
| Aggrecan (mouse ACAN core protein) | Anti-ACAN | Rabbit polyclonal<br>IgG | 500 µg/ml | Millipore<br>#AB1031 | WB<sup>2</sup> (Lendvai *et al.*, 2013 &<br>Sutkus *et al.*, 2014) |
| N-acetylgalactosamine (GalNAc) | Biotinylated Wisteria<br>floribunda agglutinin (WFA) | N/A | 2 mg/ml | Sigma<br>#L1766 | Koppe *et al.*, 1996 |
| Hyaluronan (HA) | Biotinylated Hyaluronan-<br>binding protein (Bio-HABP) | N/A | 250 µg/ml | AMSBio<br>#AMS.HKD-B141 | Tengblad, 1981 [63] |
| **Cell markers** |  |  |  |  |  |
| DAPI |  |  | - |  | - |
| Nuclear counterstain (dsDNA) | Hoescht 33342 | N/A | 10 mg/ml | Invitrogen<br>#H3570 | - |
| Neuron-specific nuclear protein (NeuN) | Anti-NeuN | Mouse<br>monoclonal IgG1 | 1 mg/ml | Millipore<br>#MAB377 | WB<sup>2</sup> (Jin *et al.*, 2003) |

**Table 2.**
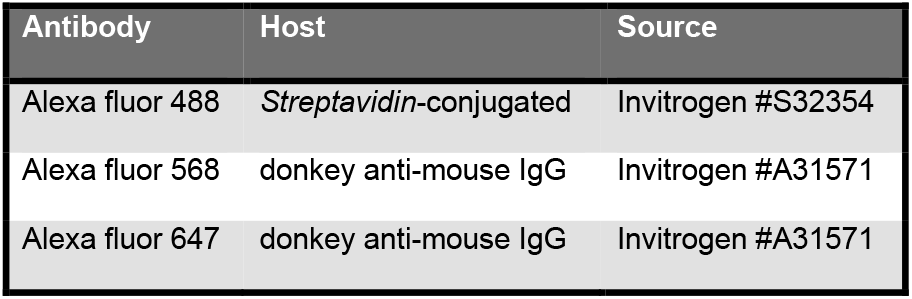
Fluorescent-conjugated secondary antibodies (2 mg/ml) used for immuno-detection of primary antibodies.

To remove primary antibodies, sections were washed again using TBS (3 × 10 mins; RT). To visualise each primary antibody staining, the tissue was then co-incubated in darkness with the fluorescent-conjugated secondary antibodies (1:500; 2 hrs; RT; Table 2) against the species of the primary antibodies. In darkness, tissues were washed in TBS (3 × 10 mins; RT) then a finally in Tris non-saline (TNS; 0.5 M Tris, pH 7.6) to reduce precipitation before air-drying. Tissues were mounted on Superfrost Plus slides, air-dried and coverslipped with FluorSave™ Reagent (EMD Millipore).

##### Image acquisition and quantification techniques

The fluorophores used to label the CNS sections were visualised using a Zeiss LSM 880 (upright) confocal microscope. Images were taken as tile scans of the entire spinal cord transverse section or a 400 μm × 1500 μm area of the cortex in brain sections at 20 × magnification (1.03 μs/pixel, averaging: 4).

To quantify PNNs in a selected area, NeuN-positive cells and their co-localisation with PNNs, as labelled by ACAN, HABP or WFA, were sequentially counted using the Cell Counter plugin (Kurt de Vos; https://imagej.nih.gov/ij/plugins/cell-counter.html) in the software FIJI [64]. The intensity (mean gray value), area and integrated density were recorded using the software FIJI [64] for ACAN, HABP or WFA for the CNS region of interest to assess the effect of 4-MU administration. For all stains, the mean gray value for the corresponding secondary control sections were subtracted from each individual measure to account for non-specific background staining.

##### Glycosaminoglycan extraction

Fresh frozen left hemisphere (12 wpi) - (n=3 Lister hooded rats per group) GAG extraction

##### Statistical analysis

Data sets were analysed and graphed using OriginPro 2019 graphing and data analysis software. Prior to statistical analysis, normality of data was determined using the Shapiro-Wilk test. For multiple group statistical analysis, Bonferroni test was used for post-hoc comparisons. Figures were produced and arranged using CorelDRAW 2018 Version 20.1.0.707 (Corel Corporation, Ottawa, Canada).

## Acknowledgements

This work was sponsored by grants from the Wings for Life Foundation (WFL-UK-008-15), Medical Research Council UK (Confidence in concept MC-PC-16050 and Project grant MR/S011110/1) to JCFK; The University of Leeds 110 Years Scholarship to SFI; Center of Reconstruction Neuroscience – NEURORECON CZ.02.1.01/0.0/0.0/15_003/0000419 to LMU, PV and JCFK; and Czech Science Agency 19-10365S to PV and JCFK).

